# Implementing rapid, robust, cost-effective, patient-centred, routine genetic testing in ovarian cancer patients

**DOI:** 10.1101/044024

**Authors:** Angela George, Daniel Riddell, Sheila Seal, Sabrina Talukdar, Shazia Mahamdallie, Elise Ruark, Victoria Cloke, Ingrid Slade, Zoe Kemp, Martin Gore, Ann Strydom, Susana Banerjee, Helen Hanson, Nazneen Rahman, for the Mainstreaming Cancer Genetics (MCG) Programme

## Abstract

**Background:** Advances in DNA sequencing have made gene testing fast and affordable, but adaptation of clinical services to capitalise on this for patient benefit has been slow. Ovarian cancer exemplifies limitations of current systems and potential benefits of increased gene testing. Approximately 15% of ovarian cancer patients have a germline mutation in *BRCA1* or *BRCA2* (collectively termed ‘BRCA’) and this has substantial implications for their personal management and that of their relatives. However, in most countries implementation of BRCA testing in ovarian cancer has been inconsistent and largely unsuccessful.

**Methods:** We developed a mainstream pathway in which BRCA testing was undertaken by cancer team members after 30 minutes online training. Patients with a mutation were sent a genetic appointment with their results. Cascade testing to relatives was performed via standard clinical genetic procedures.

**Findings:** 207 women with ovarian cancer were offered gene testing through the mainstream pathway and all accepted. 33 (16%) had a BRCA mutation. The result informed management of 79% (121/154) women with active disease including 97% (32/33) women with a mutation. All mutation-positive women and ~3.5 relatives per family have been seen in genetics. Patient and clinician feedback was very positive. >95% found the pathway to be simple and effective. The pathway offers considerable reduction in time (~5-fold) and resource requirements (~13-fold) compared to the traditional genetic pathway. We estimate it would deliver £2.6M NHS cost savings per year, and would allow implementation of national testing recommendations with existing infrastructure.

**Interpretation:** Mainstream genetic testing is effective, efficient and patient-centred and offers a mechanism for large-scale implementation of BRCA gene testing in cancer patients. The principles could be applied in many other countries and to many other areas of genomic medicine.

## INTRODUCTION

Ovarian cancer is diagnosed in 225,000 women and results in 140,000 deaths worldwide each year ^1^. Approximately 10-15% of ovarian cancer is due to germline mutations in BRCA1 or BRCA2 (collectively termed ‘BRCA’), rising to 15-20% of non-mucinous ovarian cancer ^2-6^. Determining the BRCA status of ovarian cancer patients has important therapeutic and prognostic implications ^7-10^. Additionally, it provides improved cancer risk information for family members and opportunities for predictive gene testing and risk-reducing interventions. This has been shown to be a cost-effective cancer prevention strategy ^11,12^. In view of these benefits, several countries recommend all women with non-mucinous or high-grade serous ovarian cancer have access to BRCA testing ^13-15^

Despite this recommendation the implementation of BRCA testing in ovarian cancer patients has been inconsistent and largely unsuccessful, with only 15-30% of eligible patients being offered testing ^16-19^. This disappointing and unacceptably low rate is largely due to the complexities and limitations of systems through which ovarian cancer patients currently access BRCA testing. These arose through expansion of systems developed for testing unaffected individuals with a family history of cancer, but they do not optimally serve the needs of individuals with cancer ^9^. This is particularly true for women with ovarian cancer, for whom family history is recognised to be a poor triage for BRCA testing; many ovarian cancer patients with BRCA mutations do not have a strong family history of cancer ^2,5^. Moreover, the rationales and timelines for testing cancer patients, which is increasingly performed to personalise cancer treatment, differ from that of unaffected individuals. Testing access processes need to be tailored accordingly. However, most centres use the same processes to access gene testing for individuals with and without cancer ^18,20^.

Recent technological, therapeutic and societal developments now make it imperative that modernized BRCA testing access processes for ovarian cancer patients are established. The advent of next-generation sequencing has made gene testing fast and affordable, removing the technological and economic barriers that previously hindered access. The recent approval of PARP inhibitors in the treatment of relapsed ovarian cancer in BRCA mutation carriers has further highlighted the clinical importance of testing ^10,21-24^. Finally, the knowledge, awareness and expectations of women with ovarian cancer in relation to gene testing has greatly increased in recent years, with many more women wanting access to testing ^25,26^.

Here, following wide consultation, we have developed and evaluated a patient-centred, mainstream genetic testing pathway to maximize availability, utility and equity of access to BRCA testing for women with ovarian cancer.

## METHODS

### Pathway Design

To inform pathway design we undertook wide consultation with geneticists, genetic counsellors, oncologists, cancer nurse specialists, genetic laboratories and patient representative groups throughout the UK through formal consultation days and informal information gathering ^18,26^. We designed and iterated the pathway using lean methodology ^27^. The pathway was approved by the Mainstreaming Cancer Genetics (MCG) programme board and the Royal Marsden gynaecological cancer and genetic units. Review of pathway performance was undertaken every six months. Mainstream pathway v1 was used from July 2013 to May 2014. Mainstream pathway v2 was used from May 2014 to Nov 2014 and is now the standard BRCA gene testing pathway for ovarian cancer patients at the Royal Marsden Hospital (Figure 1).

**Figure 1:**
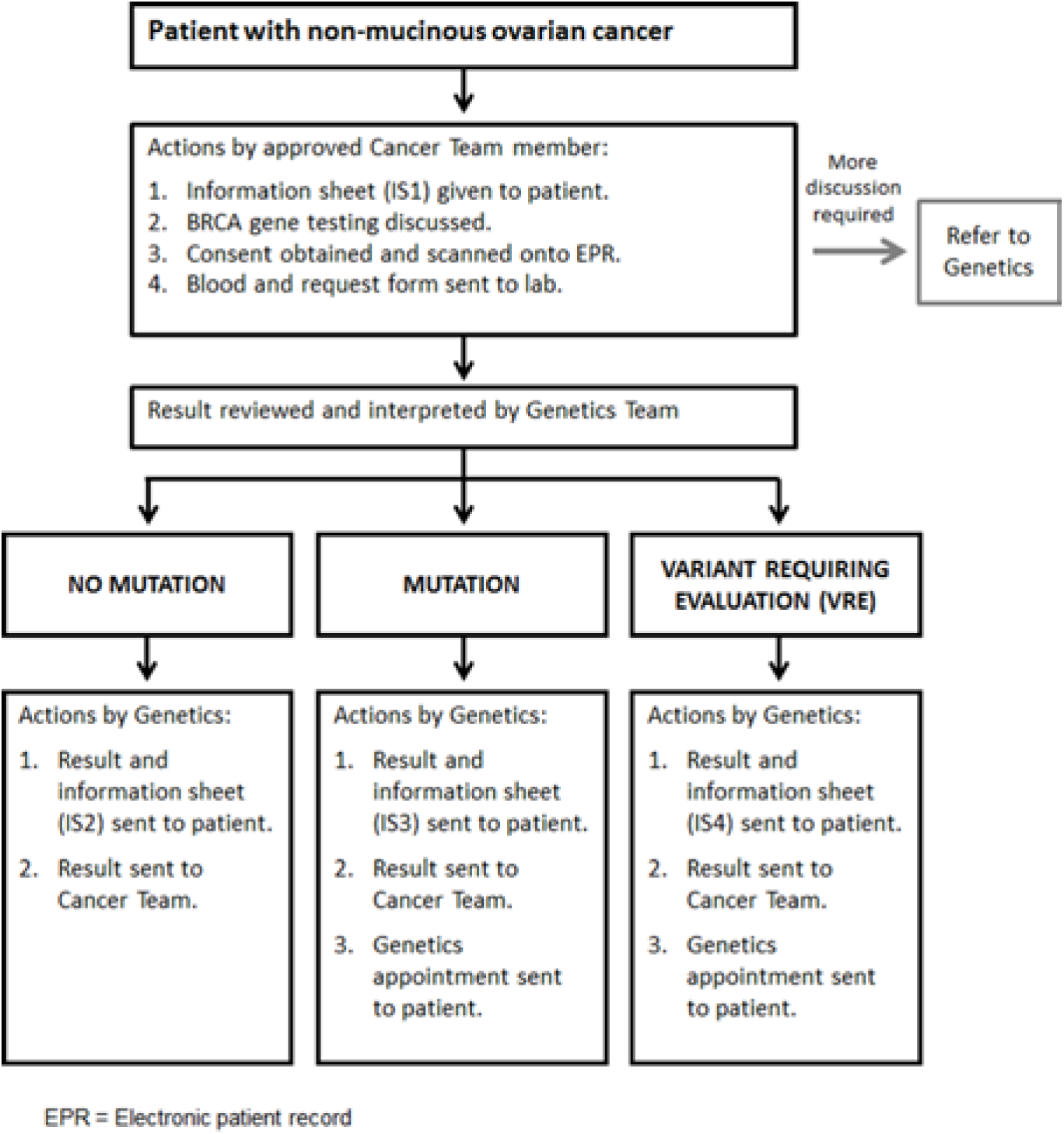
Mainstream pathway for testing in ovarian cancer patients.

### Participants

Participants were women with ovarian cancer seen between 07/2013-11/2014 who met the eligibility criteria. Between 07/2013-05/2014 the eligibility criterion was women with non-mucinous ovarian cancer below 65 years. This age limit was employed to ensure testing capacity was available. In May 2014 the age limit was removed and the testing eligibility became, and remains, any woman with non-mucinous ovarian cancer.

### Consent Training

In response to the consultation we developed a web-based e-learning training module, which can be completed in <30 minutes and is freely and easily accessible via computer, tablet or smartphone. The training includes videos, available on YouTube, and documentation about the pathway including a comprehensive frequently answered questions document. A certificate of training is provided on completion. No testing was permissible unless the training had been completed. The full training package is available on https://mcgprogramme.com/brcatesting and detailed in the Supplementary Appendix.

### BRCA gene testing

BRCA testing was performed by one of two methods: Sanger sequencing and MLPA for 127 tests before 24/03/2014, the TruSight Cancer Panel (TSCP) for 80 tests after 24/03/2014 both within the TGLclinical gene testing laboratory, which is accredited to ISO 15189 standards (www.tglclinical.com). Full details of these methods are available on request. Both methods have >99% sensitivity for small variants and exonic deletion and duplications. For both methods, pathogenic mutations were independently verified using a second aliquot of DNA by Sanger sequencing or MLPA, as appropriate. All predictive tests in relatives were also performed by Sanger or MLPA. The change in method simply reflected the evolution of TGLclinical to next-generation sequencing that occurred during the course of the study.

Interpretation of BRCA results was highlighted as a concern by both genetics and oncology ^18^. Clarity about the pathogenicity and clinical relevance of variants was considered essential. In particular, using the term ‘variant of uncertain/unknown significance’ (VUS) without further information about potential clinical management was considered confusing and liable to lead to inconsistent, inappropriate management as has been well-documented ^28,29^. Availability of all the BRCA data, rather than just select variants, was also considered desirable, but structured to ensure clinically relevant variants were clearly highlighted.

To fulfil these requirements we developed a two-page report as shown in Supplementary Figure 1. The first page is given to the patient, cancer and genetic teams and provides clear information about whether or not a pathogenic mutation was detected. The second page provides information about all the BRCA variants detected and is available on request. Pathogenic mutations were defined as variants predicted to cause premature protein truncation through frameshifting small insertions or deletions, whole exon deletion / duplications, stop-gain (nonsense) or essential splice-site changes in BRCA1 (prior to c.5545) and BRCA2 (prior to c.9924) or other types of variant (e.g. nonsynonymous (missense) variants) for which conclusive genetic proof of pathogenicity equivalent to that associated with truncating mutations was available. If a variant was plausibly pathogenic and a specific evaluation could result in definitive classification, the variant was designated a Variant Requiring Evaluation (VRE) and the specific evaluation to be performed and the time-frame for definitive classification was outlined on the report. During this study one VRE was detected, a possible exonic deletion in *BRCA2* that was likely artefactual as it was only detected by one MLPA kit. Repeat MLPA on a fresh sample confirmed that no deletion was present. All variants are also being submitted to databases to foster ongoing research. Any future reclassifications will be communicated to patients and their medical teams.

### Outcomes

The outcomes assessed were: 1) If BRCA testing was performed, 2) Result of BRCA testing, 3) Impact on patient management, 4) Impact on family management, 5) Impact on genetic service resourcing, 6) Patient experience, 7) Clinician experience.

### Cost comparison

To estimate the cost of the new pathway compared to the traditional pathway we used the pathway data and costs from Slade et al. ^20^. In the typical traditional pathway patients have a first appointment in genetics @£332.29 prior to testing and then a follow-up appointment to receive the test result @£127.38. The cost for 6,500 patients is thus £2,987,855. In the mainstream pathway the ~1000 patients with a mutation have a first appointment with genetics @£332.29 to discuss the result so the total cost of genetic appointments is £332,290. Thus the mainstream pathway has an annual cost saving of £2,655,565. All other costs e.g. for the test, follow-up of the patient and cascade to relatives, is the same in both pathways.

### Evaluation of pathway experience

We sent questionnaires to all participants six months after pathway v1 was initiated and six months after v2 was initiated to facilitate pathway optimisation and to evaluate patient experience. This was conducted through a retrospective questionnaire survey using a 5-point Likert rating scale, and was approved by the Royal Marsden Clinical Audit Committee. Questionnaires were sent to 129 patients (Supplementary Figure 2, Supplementary Table 2). The questionnaires for those with and without mutations and for those in pathway v1 vs pathway v2 were slightly different. Hence there were 4 questionnaires with 15-18 questions including 13 core questions present in every questionnaire (Supplementary Figure 2). Completed questionnaires were received from 108 patients (84% response rate), of which three had to be excluded (due to inconsistent completion) giving 89 questionnaires from women without a BRCA mutation and 16 from women with a mutation (Supplementary Table 2). Completed questionnaires were also received from 15 members of the cancer team (13 doctors and two clinical nurse specialists) that consented patients through the mainstream pathway (Supplementary Table 3).

## FINDINGS

### Mainstream genetic testing pathway

The consultation process was informative and highlighted some key themes. First, all groups wanted more access to genetic testing for ovarian cancer patients. Second, there was strong support for more flexible, patient-centred access to testing, as long as the strengths of existing systems were retained, particularly with respect to obtaining informed consent. Third, all groups believed genetic testing reports should give clear and readily understandable information about the clinical relevance of the results ^18^.

We developed a pathway in which members of the cancer team that had completed online training were able to directly consent patients to genetic testing (Figure 1, Supplementary Appendix).

In pathway v1 the result was returned to the cancer team and it was their responsibility to give the result to the patient; it was not stipulated how or when this happened. Women with a BRCA mutation were then sent a genetic appointment, ideally within 3 weeks of receiving their result. This worked well, but the six month evaluation showed room for improvement. The mutation-negative results were not consistently being communicated to patients in a timely fashion. The mutation-positive results were being communicated to patients, but the genetic team had to repeatedly check to see when this had occurred to determine when the genetic appointment letter could be sent. This was time-consuming with potential for delays and inconsistency for patients. After further consultation we revised the pathway, such that the test result was sent from genetics, in writing, to the patient and cancer team, together with the appropriate information sheet, and a genetic appointment for patients with a mutation. This ensured consistent, rapid communication of the results to all parties and all women with mutations automatically received a genetic appointment date with their result, within ~4 weeks from the test initiation. This pathway was well received by patients and the medical teams. It has become the standard pathway for BRCA testing in ovarian cancer patients in our institution. The final pathway is shown in Figure 1. Comprehensive details of the full pathway are in the Supplementary Appendix.

### Patient characteristics

207 women with ovarian cancer were offered BRCA testing through the mainstream pathway between 07/2103-11/2014 (Table 1, Supplementary Table 1).

**Table 1.** Summary of patient characteristics.

| Category | Sub-category | No mutation | BRCA1 | BRCA2 |
| --- | --- | --- | --- | --- |
| Histology | Serous | 147 | 17 | 15 |
|  | Endometrioid | 21 |  | 1 |
|  | Clear cell | 2 |  |  |
|  | Mixed epithelial | 4 |  |  |
| Age at diagnosis | Mean (range) | 57.8 (22 – 81) | 52.1 (30 – 76) | 55.9 (43 – 87) |
| Grade | Low | 5 |  |  |
|  | Intermediate | 1 |  |  |
|  | High | 168 | 17 | 16 |
| Stage | I | 23 |  |  |
|  | II | 13 | 1 | 3 |
|  | III | 107 | 14 | 11 |
|  | IV | 31 | 2 | 2 |
| Time of testing | 1st line treatment | 54 | 8 | 5 |
|  | Relapse | 70 | 6 | 3 |
|  | Remission | 42 | 3 | 6 |
|  | Remission (later relapsed) | 8 |  | 2 |
| Family history of breast cancer or ovarian cancer | No | 153 | 7 | 10 |
|  | Yes | 21 | 10 | 6 |
| BRCA status impact on treatment | No | 74 |  | 1 |
|  | Yes | 100 | 17 | 15 |

All had non-mucinous ovarian cancer; 84% (173/207) had high grade serous cancer. The average age at diagnosis was 57 years (range 22-87). 67 (32%) patients were seen during first-line treatment, 79 (38%) during treatment for disease relapse and 61(29%) during a follow-up appointment whilst in remission and off treatment, of whom 10 subsequently relapsed. All 207 women wanted testing and had testing; 200 (97%) consented to testing at first discussion. No woman requested a genetic appointment for additional discussion prior to testing. No clinician felt a woman needed additional discussions prior to testing. 37 women had a family history of a first or second degree relative with ovarian or breast cancer, of whom 31 would have been eligible for referral to genetics and testing based on 10% likelihood of having a mutation calculated according to Supplementary Figure 3.^12,30^ Full details of the participants are given in Supplementary Table 1.

### BRCA results

Pathogenic BRCA mutations were identified in 33 women (16%), 17 in BRCA1 and 16 in BRCA2 (Table 1, Figure 2, Supplementary Figure 4 and Supplementary Table 1).

**Figure 2:**
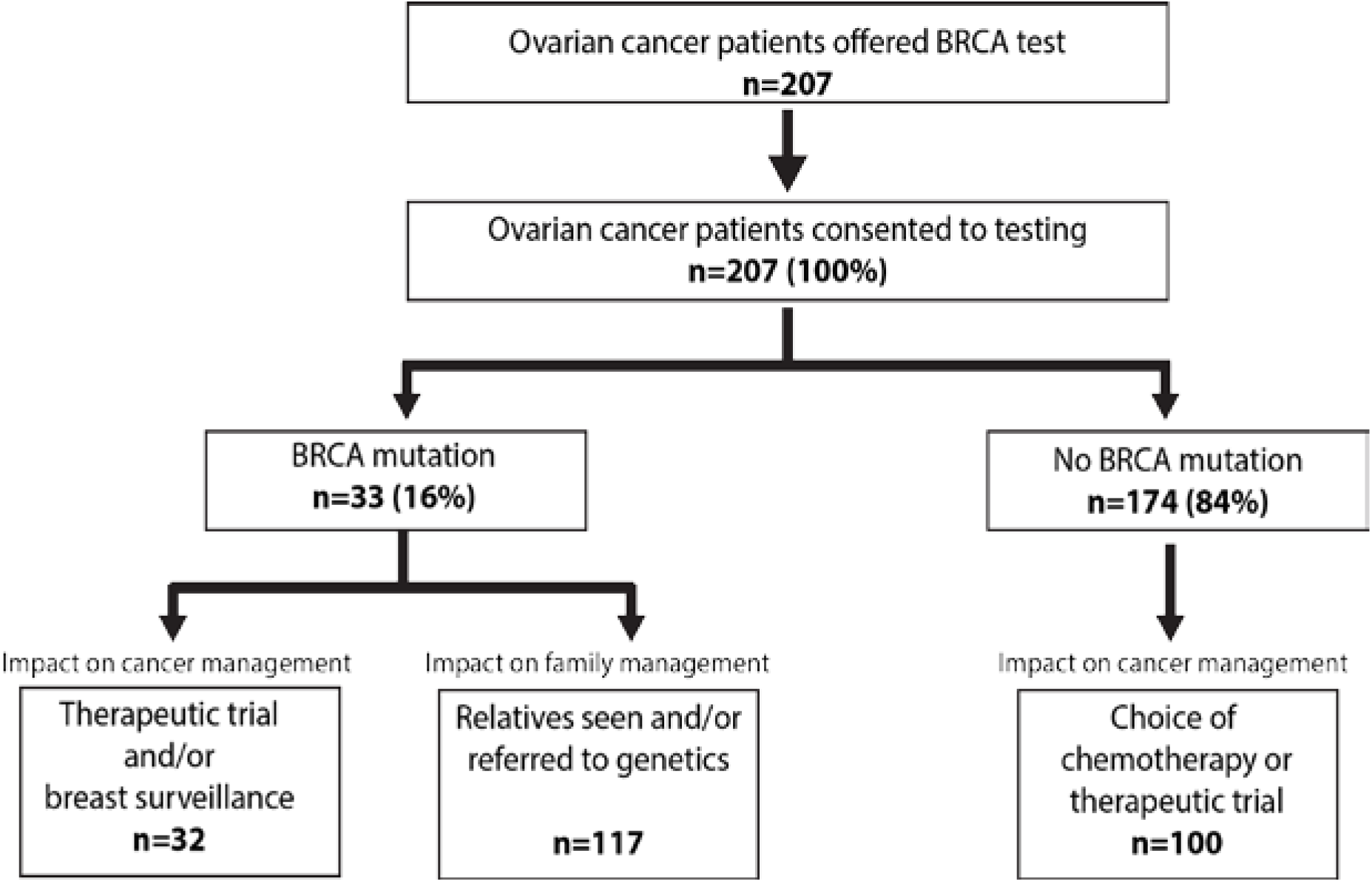
Patient flow through mainstream pathway.

Average age at diagnosis of BRCA mutation-positive individuals was 54 years (range 30-87). This was not significantly different from the mutation-negative individuals (average age at diagnosis of 58 years (range 22-79). All mutation-positive individuals had high grade cancers, 32 were serous and one was endometriod. 16 (42%) mutation-positive patients had a family history of breast or ovarian cancer in a first or second degree relative, though this had only been noted by the cancer team for 11 patients. Less than half (15/33, 45%) of the BRCA mutation-positive patients were eligible for testing based on standard genetic criteria (Supplementary Figure 3), 14 because of their family history and one because of their personal history of breast and ovarian cancer.Furthermore, only 15/39 (38%) patients in the full series eligible for testing based on genetics criteria had mutations. Five BRCA mutation-positive patients had a second cancer: two previously had breast cancer (both *BRCA1* mutation carriers), one previously had melanoma and two had synchronous

### Impact on patient management

The test result was considered useful in the management of 132/207 (64%) women, including 123/156 (79%) women with active disease (i.e. tested during first-line treatment or relapse, including women tested whilst in remission who subsequently relapsed) (Figure 2, Supplementary Table 1). For the mutation-positive individuals, the BRCA status impacted trial eligibility, with 20/23 women with BRCA mutations and active disease entered into PARP inhibitor trials, (two were too early in their disease course to be eligible for PARP inhibitor trials and one was ineligible). Furthermore, 31/32 women with mutations became eligible for enhanced breast surveillance (one woman previously had breast cancer and bilateral endometrial cancer (all *BRCA2* mutation-carriers). Thus 20% of individuals with ovarian cancer and another primary cancer had a BRCA mutation, only two of whom were eligible for testing using standard genetic criteria (Supplementary Table 1).mastectomy). Several are also having discussions with regard to possible breast cancer risk-reducing surgery. Of the 174 women without a mutation the negative result was considered in management decisions of 100/132 (75%) with active disease, primarily in relation to eligibility for trials (n=85), or choice of chemotherapy (n=15). There was no alteration in clinical management of the 42 individuals without mutations tested during remission, unsurprisingly, but the information will likely be useful for some, should they relapse in the future.

### Impact on family management

All 33 patients with a BRCA mutation attended their genetic appointment at which implications for themselves and their wider family was discussed, in accordance with standard procedures (Supplementary Figure 5). As of June 2015, 63 relatives had undergone predictive testing, of whom 33 had a BRCA mutation and 30 did not (Supplementary Figure 4). A further 54 relatives are undergoing discussions. This is an average of 3.5 relatives per family (range 0-12). No mutation-negative patient requested a genetic appointment to discuss their results further. Two mutation-negative patients were referred to genetics because of their family history of cancer.

### Patient and clinician experience

The patient experience of the mainstream pathway was very positive (Supplementary Figure 2, Supplementary Table 2). The written information was felt to be clear and helpful and no patient reported feeling unclear as to why they were offered the test. 98% felt they were given sufficient time to think about whether they wanted a test and were aware the results could have implications for themselves (96%) and their family (98%). Most patients (88%) were aware that further discussions with genetics would be organised if a mutation was found, though fewer (76%) said they were aware they could have had additional discussions with genetics prior to deciding if to have testing. Nonetheless, all were happy to have had testing and 98% were happy to have had testing at one of their cancer clinic appointments rather than at a separate appointment with genetics. All patients with mutations found their genetic appointment helpful (Supplementary Table 2). The cancer team experience was also very positive. All 15 individuals felt comfortable offering and consenting for testing and felt it was possible to consent within the time constraints of a standard oncology appointment (Supplementary Table 3).

### Large-scale implementation and cost implications

About 6,500 new patients are diagnosed with non-mucinous ovarian cancer in UK each year. All are eligible for testing based on the 10% mutation prevalence threshold recommended by NICE ^12^. We conducted a review of UK practice shortly after the NICE recommendations were announced ^18^. This revealed that a minority of ovarian cancer patients were receiving testing, primarily due to capacity limitations of genetic units together with the complexity of testing eligibility criteria, which impede referral ^20^. The traditional testing pathway requires 13,000 genetic appointments per year as all patients typically have a pre-test and post-test consultation either in person or via telephone (Figure 3).

**Figure 3:**
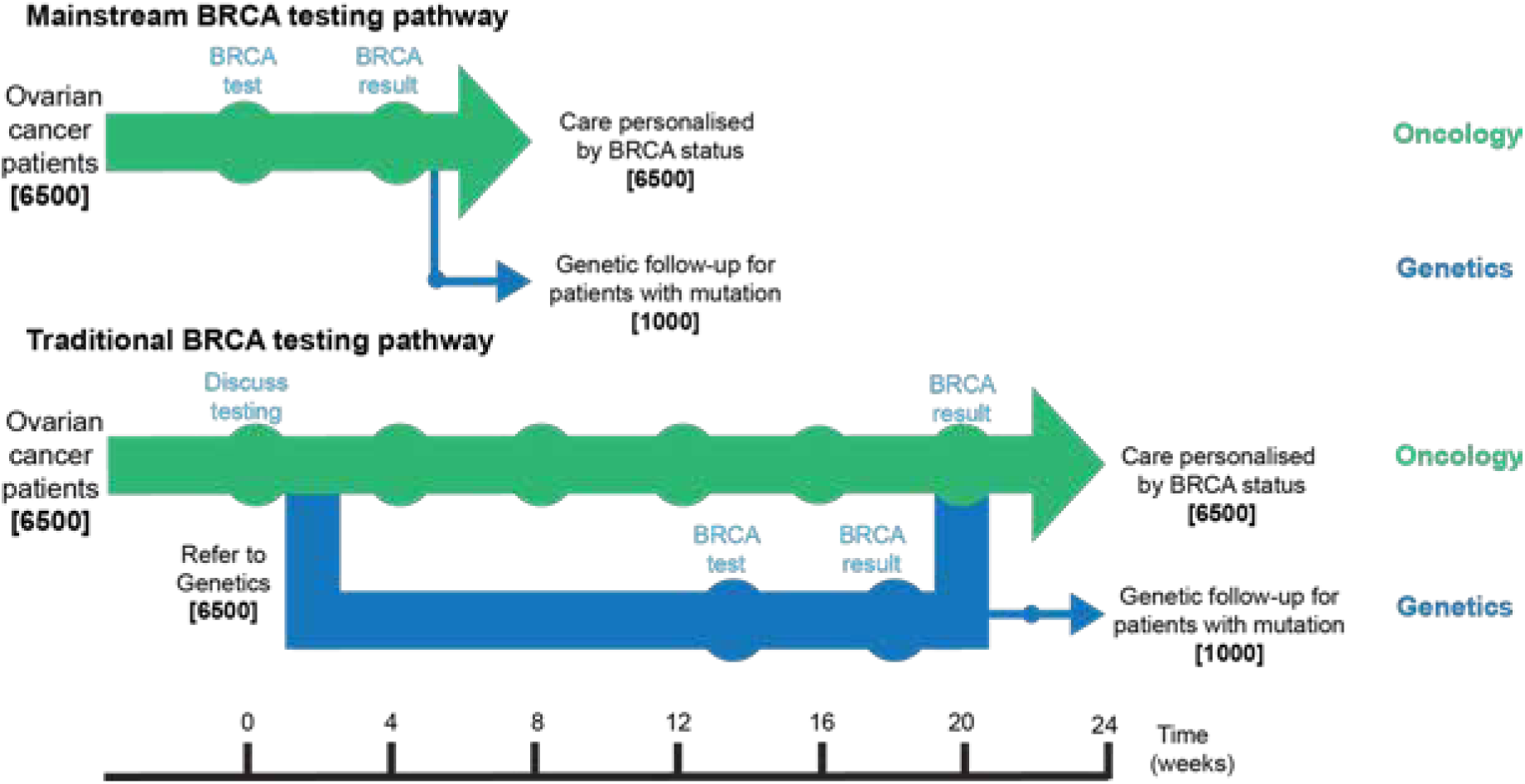
Comparison to time and appointment savings of UK-wide implementation of mainstream and traditional BRCA testing pathways. About 6,500 new patients are diagnosed with non-mucinous ovarian cancer in UK each year and are eligible for testing^12^. (a) In the mainstream pathway the testing can be completed in 3-4 weeks at existing oncology appointments and only women with a mutation (~1000) have genetic follow-up. (b) In the traditional pathway all 6,500 patients have two genetic appointments for testing leading to substantial increases in the time and resources required.

By contrast, the mainstream pathway requires ~1,000 genetic appointments (for individuals with mutations), a ~13-fold reduction, and providing a reduction in costs of ~£2.6M per year ^20^. Further cost savings would be generated in genetics as the proportion of unaffected individuals with an identified mutation in the family would increase; testing for a known mutation is 3-5x cheaper than undertaking the full gene analysis required if the family status is unknown ^20^. The mainstream pathway also offers a ~five-fold reduction in time to the result because the average waiting time for a genetic appointment in UK is 12-15 weeks (Figure 3). Use of the mainstream pathway would thus potentially allow large-scale implementation of BRCA testing in ovarian cancer patients, at a national level, with the existing clinical infrastructure.

## DISCUSSION

In this study, 16% of women with non-mucinous ovarian cancer had a BRCA mutation, adding further evidence that germline BRCA mutations are a significant cause of the disease ^2-6^. We also show that stratifying access to testing for ovarian cancer patients should be determined by personal cancer history rather than family history. Family history performs poorly as an eligibility criterion; half of the BRCA mutation carriers we identified did not have a relevant family history, and only a third of women who met the standard gene testing criteria had a mutation. Determining test eligibility on patient criteria is also much simpler to implement consistently. Moreover, the BRCA result was considered useful in management planning of 79% of patients with active disease: 97% of patients with a mutation and 75% of patients without a mutation. This is consistent with increasing data showing that stratification of ovarian cancer by BRCA status can provide important clinical, therapeutic and prognostic information, and should be part of standard care ^7,8,10^

All 207 patients offered the mainstream testing pathway accepted it. None wanted an additional genetic appointment prior to deciding whether to have a test. Furthermore, the patient feedback showed patients were pleased they had testing, irrespective of the outcome, and were pleased they could access testing through a routine cancer clinic appointment. This reflects our consultation data, and other data, which show many women with ovarian cancer want flexible, simplified access to gene testing ^25,26^. The cancer and genetic teams also had confidence in the pathway and found it straight-forward to execute.

The complexity and the time and resource (cost and personnel) requirements intrinsic to most genetic testing pathways has impeded routine access for ovarian cancer patients. The traditional pathway involves all cancer patients consulting genetics before and after testing. This hinders timely availability of results, and introduces multiple points for patients to be lost from the system before getting their result. Here, we have streamlined the pathway, without compromise of quality or patient care, by training the cancer team to provide information for patients to decide if they want to have a test. This allows genetic expertise to be concentrated where it is needed, namely the patients with mutations and anyone with questions that cannot be addressed by the cancer team. It is also convenient and appropriate in an era of increasing tumour genetic tests being ordered and consented for by cancer teams. Tumour testing can reveal variants that are germline cancer predisposition mutations. It is thus imperative that the consenting and patient management processes for germline and tumour testing are integrated. Additionally, the mainstream pathway greatly reduces the time, appointments and cost of gene testing compared to the conventional pathway. We estimate the reduction in pre-test genetic consultations alone would save the NHS >£2.6M annually.

Change to a mainstream testing pathway will likely lead to some new challenges, at least in the short-term. For example, in the set-up phase, in addition to new ovarian cancer cases, surviving women with a previous diagnosis of ovarian cancer will also be eligible for testing. In the UK, there are ~25,000 such women for whom test resourcing will need to be secured. A longer term challenge relates to accuracy of risk information provided to unaffected relatives of BRCA mutation-carriers identified through ovarian cancer patient testing. The cancer risk figures typically used are derived from multi-case high-risk families. These can over-estimate the risks to relatives of mutation carriers identified in different contexts, for example testing of cancer patients, or as incidental findings ^31,32^. Reliable risk estimates in these other contexts are currently not available, and are urgently needed. That being said, given that no effective screening for ovarian cancer is available, the disease often presents at advanced stage and mortality is still considerable, it is likely that salpingectomy and oophorectomy, which reduces ovarian cancer risk by >80%, will be an attractive option for many unaffected BRCA mutation-carriers identified through mainstream testing of ovarian cancer patients ^33,34^. This is also likely to be cost-effective for healthcare systems given that existing data indicate the much more expensive traditional BRCA testing pathway is cost-effective at a 10% mutation detection threshold ^11,12^. Of equal importance, it would expedite prevention of the thousands of ovarian cancers that occur due to germline BRCA mutations.

The mainstream gene testing pathway we describe here could be readily implemented, at a national level, in the UK and other countries. It is an effective, efficient, patient-centred pathway that allows genetic testing access for ovarian cancer patients to be increased in a consistent and equitable fashion. We have initiated a similar study for BRCA testing in breast cancer patients and are currently expanding the approach to accommodate gene panels. The principles could equally be applied to other cancer predisposition genes and other areas of genomic medicine ^9,32,35^. To facilitate this we have made the components of our pathway fully and freely available in the Supplementary Appendix and on the programme’s website: www.mcgprogramme.com

## Acknowledgements

We thank the patients and clinicians of the Gynaecological Cancer Unit and Cancer Genetics Unit at The Royal Marsden for their participation. We thank the many people that participated in our consultation events for their time and engagement. We acknowledge NHS funding to the Royal Marsden/ICR NIHR Specialist Biomedical Research Centre for Cancer. This study was overseen by the MCG programme committee and collaborators. See www.mcgprogramme.com for full details of individuals involved in the MCG programme.

## Authors’ contributions

NR, AG, HH, MG and SB designed the oncogenetic pathway. AG, MG and SB led implementation in the oncology clinical services. HH, AG and NR led implementation in the clinical genetics service. SS, SM, VC, ER and AS performed BRCA testing and analyses. DR, AG, NR, IS, HH, ZK, ST and AS designed and analysed the patient and clinician evaluation. NR and AG wrote the manuscript with contributions from all the other authors. DR project managed the study. NR oversaw the study.

